# Identification of *Pappa* and *Sall3* as Gli3 direct target genes acting downstream of cilia signalling in corticogenesis

**DOI:** 10.1101/2024.04.16.589766

**Authors:** Shinjini Basu, Lena Mautner, Kae Whiting, Kerstin Hasenpusch-Theil, Malgorzata Borkowska, Thomas Theil

**Author notes:** equal contributors (Shinjini Basu, Lena Mautner, Kae Whiting).

## Abstract

The cerebral cortex is critical for advanced cognitive functions and relies on a vast network of neurons to carry out its highly intricate neural tasks. Generating cortical neurons in accurate numbers hinges on cell signalling orchestrated by primary cilia to coordinate the proliferation and differentiation of cortical stem cells. While recent research has shed light on multiple ciliary roles in corticogenesis, specific mechanisms downstream of cilia signalling remain largely unexplored. We previously showed that an excess of early-born cortical neurons in mice mutant for the ciliary gene *Inpp5e* was rescued by re-introducing Gli3 repressor. By comparing expression profiles between *Inpp5e* and *Gli3* mutants, we here identified novel Gli3 target genes. This approach highlighted the transcription factor gene *Sall3* and *Pappalysin1* (*Pappa*), a metalloproteinase involved in IGF signalling, as up-regulated genes. Further examination revealed that Gli3 directly binds to *Sall3* and *Pappa* enhancers and suppresses their activity in the dorsal telencephalon. Collectively, our analyses provide important mechanistic insights into how primary cilia govern the behaviour of neural stem cells, ultimately ensuring the production of adequate numbers of neurons during corticogenesis.

**SUMMARY STATEMENT:** This study reports how cilia control gene expression via Gli3 in the developing murine cerebral cortex.

## INTRODUCTION

The cerebral cortex consists of dozens of different types of neurons to perform highly complex neural tasks (van den Ameele et al., 2014). Understanding how these neurons are generated in correct quantities, at the right time and place poses a major challenge. Corticogenesis entails a multi-step process beginning with the subdivision of the telencephalon into distinct dorsal and ventral domains that give rise to the neocortex and the basal ganglia, respectively. This patterning process coincides with an expansion of cortical stem and progenitor cells that eventually undergo neurogenesis to form the various neuronal subtypes in a coordinated manner. These processes heavily rely on extensive cell signalling facilitated by primary cilia, tiny cell surface protrusions that act as antennae for cell signals. Cilia are critically important for controlling cortical growth in mice (Foerster et al., 2017; Wilson et al., 2012) and in humans (Bachmann-Gagescu et al., 2012; Budny et al., 2006; Davis et al., 2007; Jamsheer et al., 2012; Putoux et al., 2011) and regulate the activity of signalling pathways essential for cortical progenitor development (Foerster et al., 2017; Wilson et al., 2012). Notably, they govern the formation of the Gli3 repressor (Gli3R) crucial for cortical growth (Hasenpusch-Theil et al., 2020; Hasenpusch-Theil et al., 2018; Wang et al., 2011; Wang et al., 2014; Wilson et al., 2012). These findings strongly support a role for cilia in controlling cortical stem cell behaviour, but the underlying mechanisms have hardly been investigated.

We recently addressed this question by analysing mice with a mutation in the *Inositol Polyphosphate-5-Phosphatase E* (*Inpp5e*) gene which regulates the phosphoinositol composition of the cilium and thereby ciliary protein trafficking and signalling (Chavez et al., 2015; Constable et al., 2020; Garcia-Gonzalo et al., 2015; Hasenpusch-Theil et al., 2020). The analysis of *Inpp5e* mutants unveiled a profound role of cilia in cortical stem cells since mutant radial glial cells (RGCs) predominately underwent direct neurogenesis resulting in increased deep layer neuron formation (Hasenpusch-Theil et al., 2020). This phenotype coincided with reduced Gli3R formation and was remarkably rescued by re-introducing Gli3R. Additionally, human cortical organoids lacking *INPP5E* function were ventralized due to reduced GLI3R levels and increased SHH signalling (Schembs et al., 2022). These findings indicate an evolutionarily conserved role of *INPP5E* in controlling GLI3R formation during corticogenesis but the downstream genes and processes remained unclear.

Here, we systematically analysed cortical development in Inpp5e mutant mice using gene expression profiling. A comparison with an mRNA-seq data set from *Gli3* conditional mouse mutants (Hasenpusch-Theil et al., 2018) revealed a significant overlap in differentially expressed genes (DEGs) suggesting a convergence onto a common phenotype. As Gli3 primarily acts as a transcriptional repressor during corticogenesis (Fotaki et al., 2006), we focussed on a common set of up-regulated genes involved in dorsal/ventral patterning, cilia disassembly and known Sonic hedgehog target genes. *Pappalysin* (*Pappa*), a regulator of insulin growth factor (IGF) signalling (Lawrence et al., 1999), and the transcription factor gene *Spalt-like 3* (*Sall3*) were amongst the most strongly up-regulated genes and were ectopically expressed in the mutant dorsal telencephalon. Furthermore, Gli3 protein bound to *Pappa* and *Sall3* enhancers, and mutations in these Gli3 binding sites led to ectopic enhancer activity in cortical progenitors. These findings establish *Pappa* and *Sall3* as novel Gli3 target genes and suggest their involvement downstream of cilia signalling and Gli3R in controlling cortical neurogenesis.

## RESULTS AND DISCUSSION

### Gene expression profiling of *Inpp5e* mutant dorsal telencephalon

We recently reported that the *Inpp5e* mutation alters the balance between direct and indirect neurogenesis (Hasenpusch-Theil et al., 2020). To explore broader gene expression changes underlying this phenotype, we performed bulk mRNA sequencing (mRNA-seq) to compare the expression profiles in the dorsal telencephalon of E12.5 control and *Inpp5e* mutant embryos. This analysis identified 2533 DEGs (padj < 0.05), with 1329 up-regulated and 1204 down-regulated genes (Table S1). Gene ontology (GO) analysis showed that these genes were primarily involved in neuronal differentiation (GO:BP terms: “regulation of neuron differentiation”, “axonogenesis”, “synapse organisation”, “regulation of membrane potential”) and forebrain development (Fig. 1A). Down-regulated genes related to “forebrain development” and “negative regulation of neurogenesis” whereas up-regulated genes were associated with “positive regulation of neurogenesis”, “axon guidance” and “regulation of membrane potential” (Fig. 1B, C). These categorisations aligned with our previous observations of mild patterning, neurogenesis and axon pathfinding defects in *Inpp5e* mutants (Hasenpusch-Theil et al., 2020; Magnani et al., 2015). To better understand how *Inpp5e* regulates these processes, we created a network plot of gene connections, highlighting *Fgfr3, Hairy and enhancer of split 1* (*Hes1*) and *Inhibitor of DNA binding 4* (*Id4*) among the down-regulated genes (Fig. S1). These genes are involved in Fgf, Notch and Bmp signalling, respectively, suggesting alterations in these pathways may contribute to the partial ventralisation and increased neurogenesis. Accordingly, we observed up-regulation of key regulators of ventral telencephalon development and down-regulation of genes governing dorsal telencephalon development (Fig. 1D). Taken together, these findings support *Inpp5e*’s previously described roles in forebrain patterning and neuronal differentiation.

**Figure 1:**
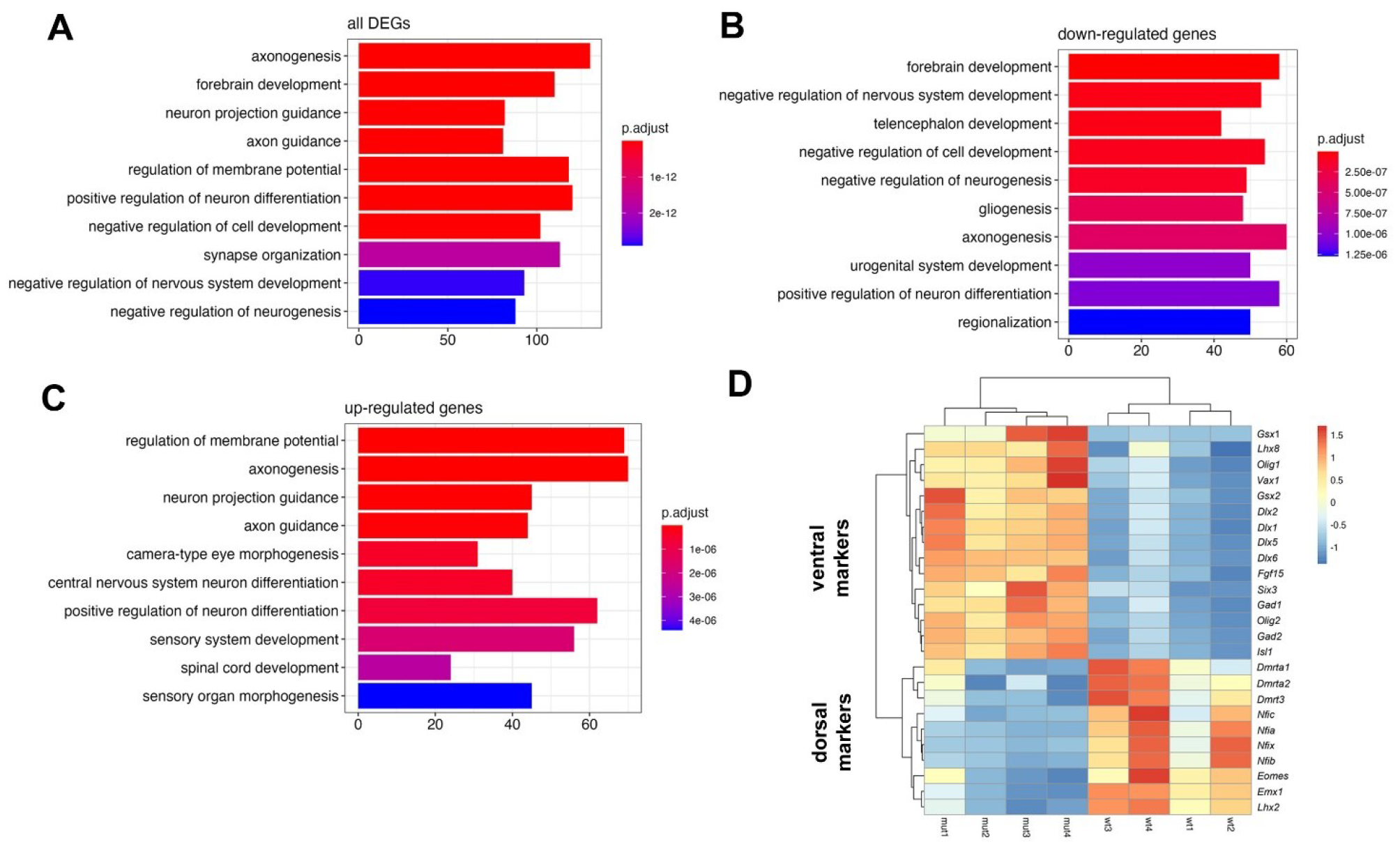
Differential gene expression in the E12.5 *Inpp5e*^Δ/Δ^ dorsal telencephalon. (A-C) Gene ontology (GO) analysis of all differentially expressed genes (A), and of only down-regulated (B) or up-regulated genes (C). (D) Heatmap comparing the expression of dorsal and ventral telencephalic markers.

### Identification of genes acting downstream of *Gli3* in *Inpp5e* mutants

We previously reported that re-introducing Gli3R in *Inpp5e* mutants rescued the imbalance between direct and indirect neurogenesis (Hasenpusch-Theil et al., 2020). To identify downstream genes of *Gli3* that potentially mediate this rescue, we compared the *Inpp5e* gene expression profiling with our bulk mRNAseq analyses of dorsomedial telencephalon of E11.5 and E12.5 *Gli3* conditional mouse mutants in which *Gli3* is inactivated using an *Emx1Cre* driver line (Hasenpusch-Theil et al., 2018). Genes differentially expressed in both mutants are candidates to be regulated by *Gli3* in *Inpp5e* mutants. This comparison revealed statistically significant overlaps in DEGs between *Inpp5e* and *Gli3* mutants at E11.5 and E12.5 (Fig. 2A), (Table S1) with nearly 50% of all DEGs in E12.5 *Gli3* mutants differentially expressed in *Inpp5e* embryos. We also observed correlations between the fold changes in the two mutants (Fig. 2B, C). While 24% of genes were regulated oppositely at E11.5, a remarkable 95% of DEGs were either up-regulated or down-regulated at E12.5 suggesting a convergence of phenotypes despite some differences in the dissected tissue and in the effects of the mutations on Gli3R. Whereas *Inpp5e* mutants showed increased neuron formation in the dorsolateral telencephalon, E11.5 *Gli3* conditional mutants initially exhibited delayed neurogenesis in the rostromedial dorsal telencephalon which resolved by E12.5. This discrepancy likely stems from variations in the analysed tissues and reflects the lateral to medial neurogenic gradient. Notably, negative regulators of neurogenesis such as *Pleiotrophin* (*Ptn*) and *Mycn* were down-regulated in *Inpp5e* embryos but up-regulated in E11.5 *Gli3* conditional mutants (Table S1).

**Figure 2:**
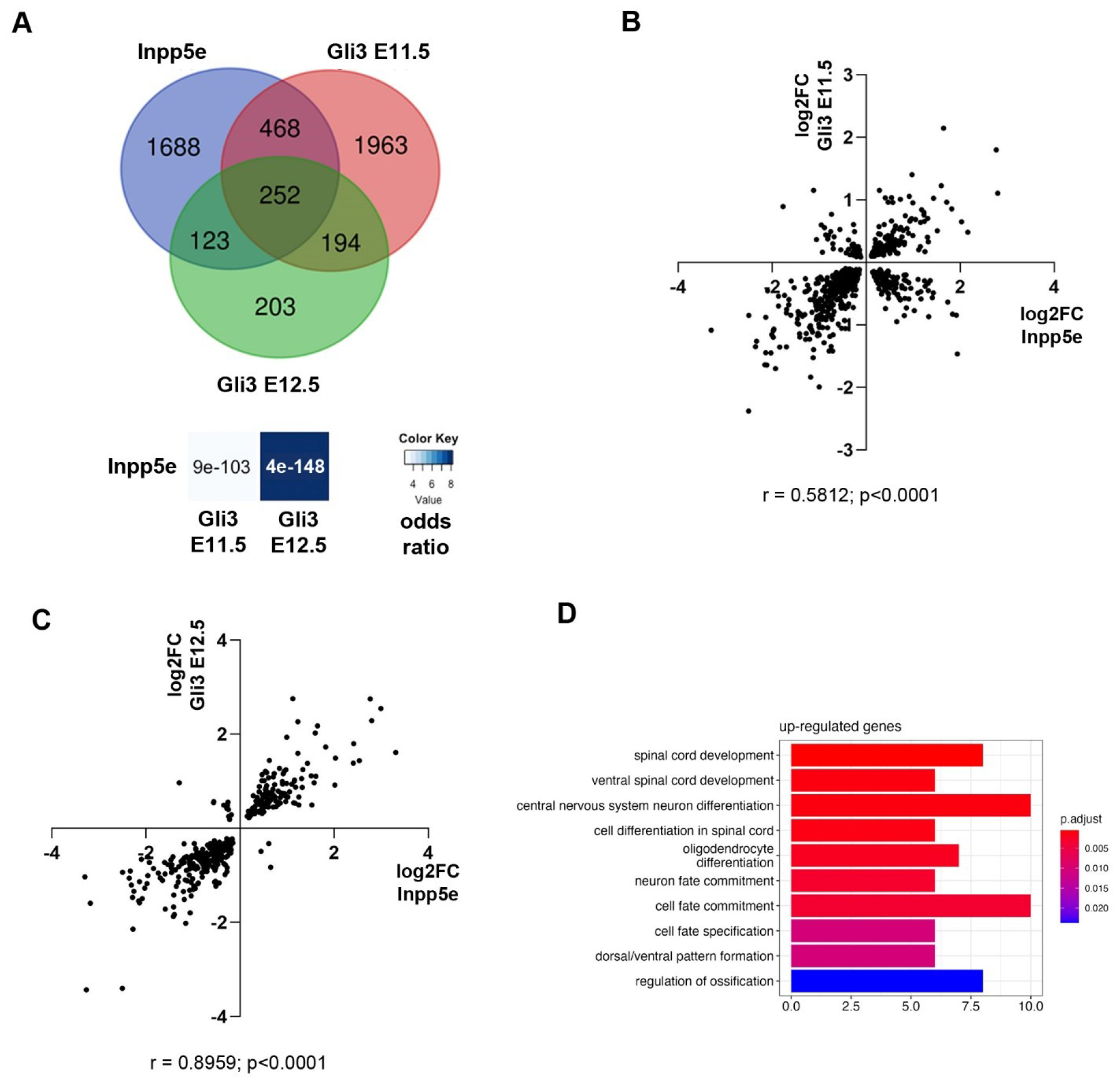
Comparison of differential gene expression between *Inpp5e* and *Gli3* mutants. (A) Venn diagram intersection of DEGS in E12.5 *Inpp5e*^Δ/Δ^, E11.5 and E12.5 *Gli3*^cKO^ embryos. Significance and odds ratio are indicated. (B, C) Comparison of gene expression changes between *Inpp5e*^Δ/Δ^ and E11.5 (B) and E12.5 (C) *Gli3*^cKO^ mutants. (D) GO analysis of genes up-regulated in both mutants. Statistical tests: Fisher’s exact test (A) and Spearman correlation (B, C).

As *Gli3* predominately acts as a repressor in cortical development (Fotaki et al., 2006), we focussed on the 135 genes that were up-regulated in both mutants at E12.5. Among the top six up-regulated genes are *Gsx2, Olig1/2*, and *Fgf15*, critical for ventral telencephalic development. We also noted an increased expression of the Shh target genes *Patched1* (*Ptch1*) and *Cyclin D1* (*Ccnd1*), emphasized by GO:BP terms such as “Dorsal/ventral pattern formation” and “oligodendrocyte differentiation” (Fig. 2D). Closer inspection also revealed an up-regulation of *Histone deacetylase 6* (*Hdac6*), a regulator of ciliary disassembly (Forcioli-Conti et al., 2016; Loktev et al., 2008; Lysyganicz et al., 2021; Pugacheva et al., 2007), suggesting a novel feedback mechanism whereby cilia mediated Shh signalling stimulates *Hdac6* expression which in turn may lead to more labile or shorter cilia with reduced signalling capacity (Macarelli et al., 2023). Overall, these gene expression changes align with the partial ventralisation of the dorsal telencephalon in both mutants and the destabilised cilia in *Inpp5e* mutant embryos.

### *Pappa* and *Sall3* expression are elevated in *Gli3* conditional and *Inpp5e* mutants

The strong overlap of DEGs in *Inpp5e* and *Gli3* mutants provided us with a unique foundation for identifying genes acting downstream of cilia and Gli3. To explore this, we first performed *in situ* hybridisations focussing on genes with relevance to telencephalic patterning and cell signalling. Notably, *Sall3* and *Pappa* were amongst the most strongly up-regulated genes and encode a zinc finger transcription factor and a zinc metalloproteinase involved in IGF signalling (Lawrence et al., 1999), respectively. Our expression analysis revealed that in control embryos *Pappa* and *Sall3* transcripts were confined to ventral telencephalic progenitors but were found throughout the rostral cortex of *Gli3* mutants (Figure S2). In *Inpp5e* mutants, the effect was less pronounced and their transcription extended into the dorsolateral telencephalon. These patterns confirmed the up-regulation of both genes and validated our bulk mRNA-seq results. The varying degrees of ectopic expression likely result from different impacts of the mutations on Gli3R levels which is decreased by approximately 50% in *Inpp5e* mutants (Hasenpusch-Theil et al., 2020), whereas Gli3R is completely lost in *Gli3* conditional embryos from E11.5 (Hasenpusch-Theil et al., 2018).

### Gli3 binds to *Pappa* and *Sall3* forebrain enhancers *in vivo* and *in vitro*

The up-regulation of *Pappa* and *Sall3* suggested that Gli3 may directly control their expression by binding to and repressing gene regulatory elements in cortical cells, thereby restricting their transcription to the ventral telencephalon. To test this hypothesis, we examined a Gli3 ChIP-seq data set (Hasenpusch-Theil et al., 2018) and identified a Gli3 peak within the *Pappa* gene overlapping with exon 13 and coinciding with a region of open chromatin in E11.5 forebrain tissue (ENCODE accession number ENCFF426VDN) (Fig. 3A). A 1 kb sequence surrounding exon 13 contained three potential Gli3 binding sites. Site 1 located within exon 13 and showed high evolutionary conservation across several vertebrate species while sites 2 and 3 within intron 12 were less conserved (Fig. 3C). Importantly, the human site 3 contained an A/T exchange in a critical nucleotide of the Gli3 binding sequence and a GLI3 Cut&Tag experiment on human cortical organoids showed a GLI3 peak only encompassing sites 1 and 2, but not site 3 (Fleck et al., 2022). Hence, we focussed our further analyses on sites 1 and 2.

**Figure 3:**
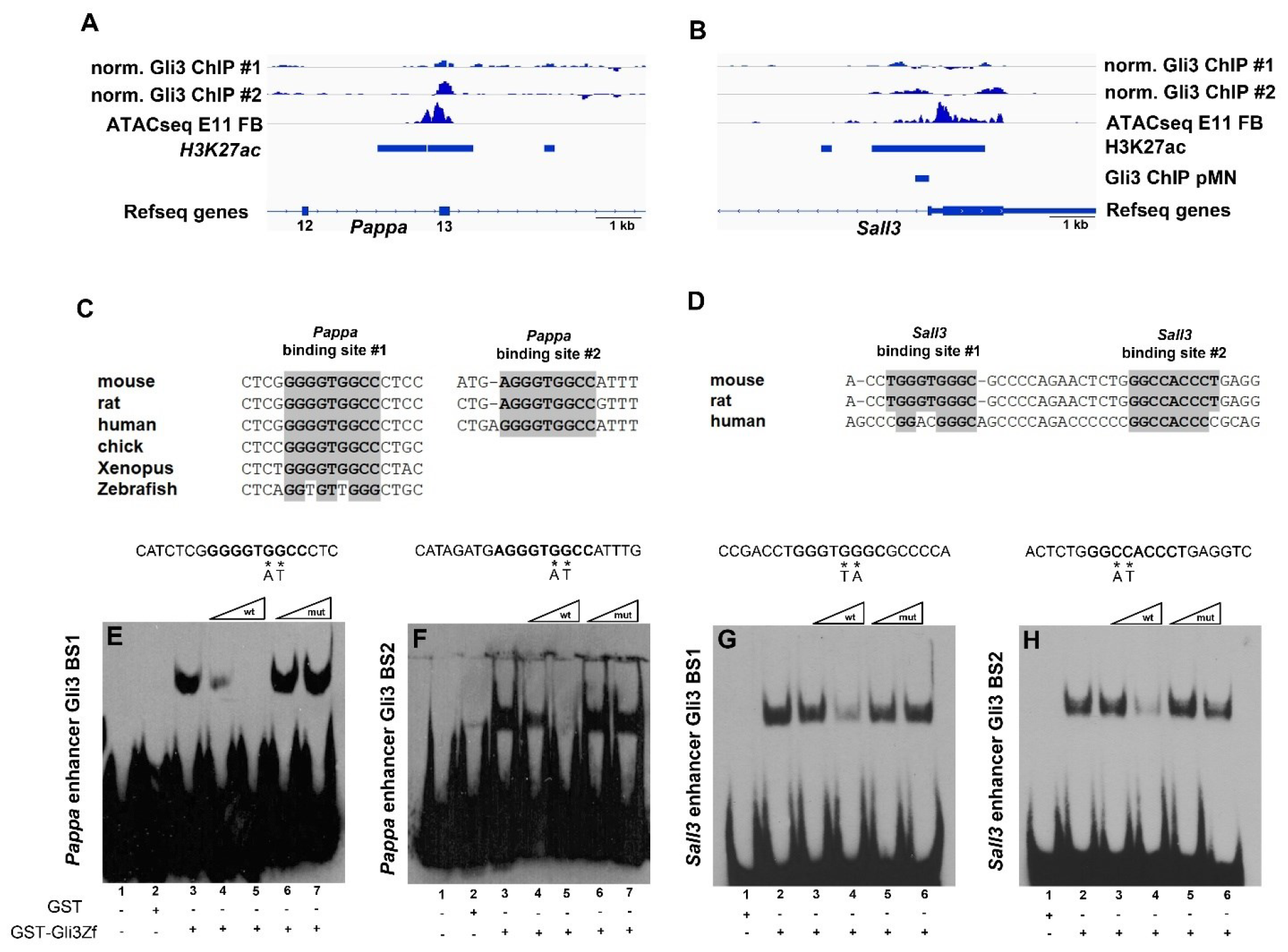
Gli3 binds to *Pappa* and *Sall3* enhancers in vivo and in vitro. (A, B) Genome browser snapshots showing Gli3 ChIP-peaks at exon 13 of *Pappa* overlapping with an open chromatin region (ATACseq peak) (A) and in the first intron of *Sall3* (B). The latter coincides with an H3K27ac positive region and a Gli3 ChIP-peak identified in motor neuron progenitors. (C, D) Evolutionary conservation of Gli3 binding sites in the *Pappa* (B) and *Sall3* (D) enhancers. (E-H) EMSAs demonstrating specific binding of a GST-Gli3 fusion protein to binding sites 1 (E) and 2 (F) of the *Pappa* enhancer and to sites 1 (G) and 2 (H) of the *Sall3* enhancer.

For *Sall3*, we noted several Gli3 peaks surrounding the transcriptional start site (Fig. 3B). The intronic peak overlapped with an open chromatin region and with Gli3 binding peaks in murine motor neuron progenitors (Nishi et al., 2015) and human cortical organoids (Fleck et al., 2022). This region contained two adjacent Gli3 binding sites in mouse and rat whereas only one site was present in the human genome (Fig. 3D).

To confirm Gli3 binding to these sites *in vitro*, we utilized a GST-Gli3 fusion protein containing the Gli3 DNA binding domain in electromobility shift assays (EMSAs) with biotin-labelled oligonucleotides encompassing the binding motifs from the *Pappa* and the *Sall3* genes. This approach resulted in the formation of a slower migrating complex for all binding sites (Fig. 3E-H). Competition assays using un-labelled wild-type oligonucleotide progressively reduced binding with increasing amounts of the competitor, while oligonucleotides with a GG to AT exchange, abolishing Gli binding (Hepker et al., 1999), did not affect complex formation. Thus, Gli3 specifically bound to sequences within the *Pappa* and *Sall3* genes.

### Gli3 represses *Pappa* and *Sall3* forebrain enhancer activity

Finally, we assessed the *in vivo* functionality of the Gli3 binding sites. We subcloned wild-type or *Gli3* binding motif mutant *Pappa* and *Sall3* enhancers into the pGZ40 reporter vector containing a *lacZ* reporter gene under the control of a human *β-globin* minimal promoter. These reporter gene constructs were co-electroporated with a GFP expression plasmid into the forebrain of E13.5 embryos which were harvested 24 hours post-electroporation. Adjacent cryosections were subsequently stained with X-Gal and a GFP antibody to monitor enhancer activity and reveal transfected cells, respectively (Fig. 4). Despite extensive electroporation, the wild-type *Pappa* enhancer only exhibited mild activity in the dorsolateral telencephalon after 24 hours of staining (Fig. 4A, B), consistent with *Pappa* gene expression being confined to the ventral telencephalon. In contrast, the mutant enhancer construct elicited strong enhancer activity in dorsolateral cortical stem cells only after three hours of staining (Fig. 4F, G). Similarly, the wild-type *Sall3* enhancer led to weak β-galactosidase staining in very few cells immediately dorsal to the pallial-subpallial boundary in three out of five electroporated brains (Figure 4C-E). Embryos electroporated with the mutant *Sall3* enhancer showed many, strongly stained cells in an extended region in three out of four embryos (Figure 4H-J). These findings suggest that the Gli3 binding sites are essential elements in repressing *Pappa* and *Sall3* expression in the dorsal telencephalon.

**Figure 4:**
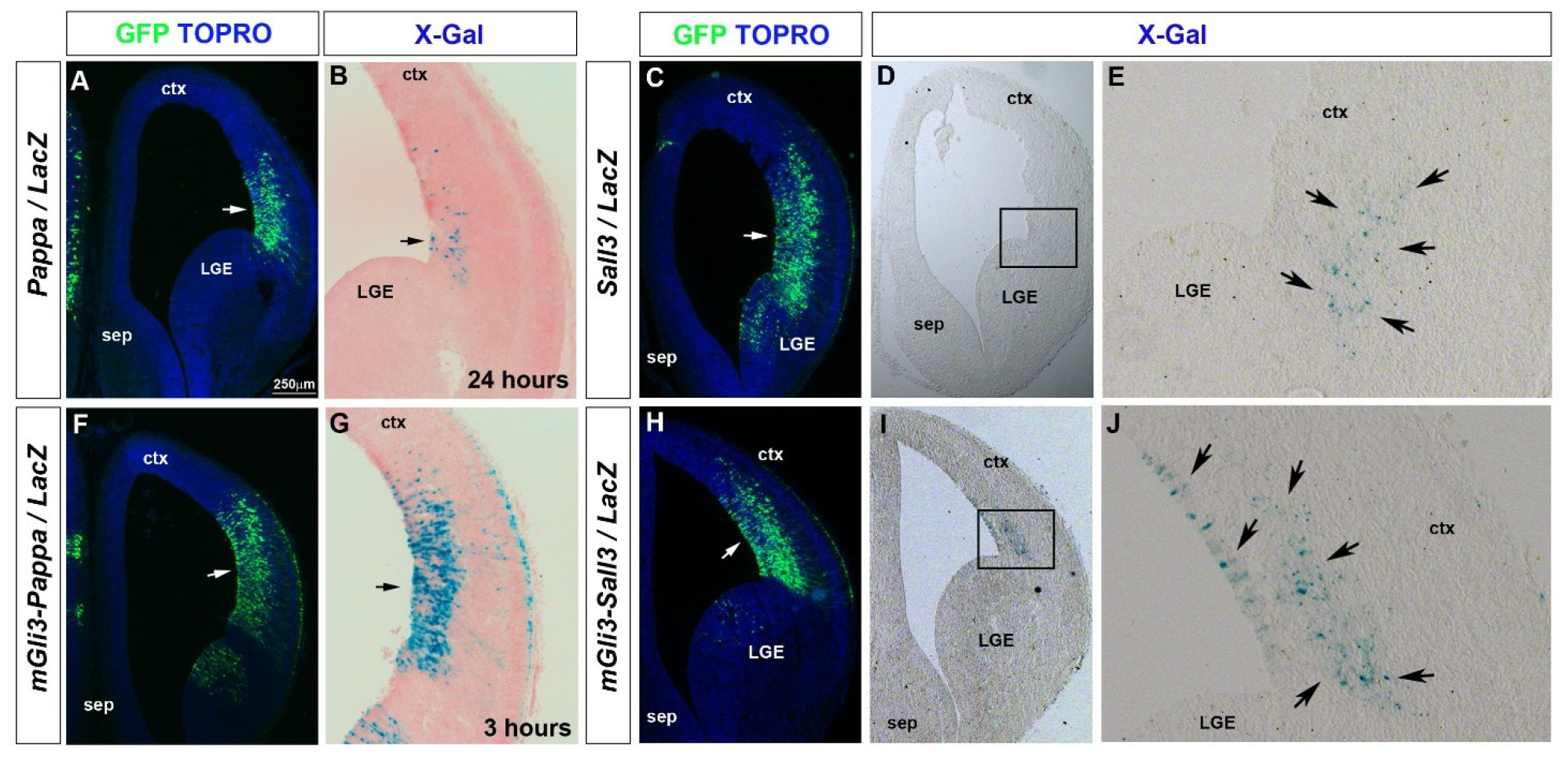
Gli3 represses activity of *Pappa* and *Sall3* enhancers in the dorsolateral telencephalon. Coronal forebrain sections of E14.5 embryos in utero electroporated with the indicated constructs were stained either with GFP antibodies or with X-Gal. GFP staining indicates the electroporated regions (white arrows) (A-E) The *Pappa* and *Sall3* enhancers showed weak activity in a limited number of cells in the dorsolateral telencephalon. (F, G) Mutations in the Gli3 binding sites led to strong reporter gene expression (black arrows). Note the different staining times. (H-J) Activity of a Gli3 binding site mutant *Sall3* enhancer was stronger and more widespread (black arrows). Abbreviations: ctx, cortex; MGE, medial ganglionic eminence; LGE, lateral ganglionic eminence; sep, septum. Scale bar: 250μm.

Taken together, the results from our expression analyses, DNA binding and reporter gene essays establish *Pappa* and *Sall3* as novel direct Gli3 targets. Their roles in corticogenesis have not been studied, but their known functions in other developmental contexts offer intriguing possibilities. Previous studies have shown complex interactions between *Sall* genes and the *Gli3*/*Shh* pathway and placed these genes upstream (Kawakami et al., 2009) and downstream of Shh signalling (Kawakami et al., 2009; Nishi et al., 2015) or revealed cooperative interactions (Akiyama et al., 2015). *Sall* gene function in neural development is only poorly understood and is complicated by complex and overlapping expression patterns (Supp. Fig. 3) (Bohm et al., 2008; Harrison et al., 2012; Ott et al., 2001; Sato et al., 2003) suggesting potential redundant functions as seen during limb development and neural tube closure (Bohm et al., 2008; Kawakami et al., 2009). *SALL3* deletion is associated with 18q23 deletion syndrome, characterized by intellectual disability and limb abnormalities. In mice, loss of *Sall3* resulted in palate deficiency, abnormalities in cranial nerves, and perinatal lethality (Parrish et al., 2004). While telencephalic development was not explored in this mutant, ectopic *Sall3* expression in the cortical primordium might interfere with Sall1’s nuclear transport and function as a transcription factor (Sweetman et al., 2003). This might lead to premature neuronal differentiation and increased neuron formation as observed in *Sall1* global and conditional mutants (Harrison et al., 2012).

Pappa plays a critical role in Igf signalling by proteolytically cleaving Igf binding proteins (Igfbps), thereby releasing sequestered Igfs for signalling (Lawrence et al., 1999). These secreted factors, their receptors and Igfbps are expressed in the developing cortex and surrounding tissue (Supp. Fig. 4) (Ayer-le Lievre et al., 1991; Bondy et al., 1992; Higginbotham et al., 2013) suggesting that cortical RGCs are responsive to Igfs. Interfering with Igf signalling reduced brain growth, while Igf2 from the cerebrospinal fluid stimulated neural progenitor proliferation (Beck et al., 1995; Kappeler et al., 2008; Lehtinen et al., 2011; Liu et al., 2009). Hence, *Pappa*’s widespread up-regulation likely contributes to increased proliferation in E11.5 *Gli3* conditional mutants. In contrast, the restricted ectopic *Pappa* expression in *Inpp5e* mutants coincided with an increase in direct neurogenesis. Interestingly, Igf signalling can promote neuronal differentiation under certain conditions. *Nestin*/*Igf1* transgenic mice showed a preferential increase in the formation of layer V neurons (Hodge et al., 2005) and Igf2 also promoted adult neural stem cell differentiation through upregulation of *Cdkn1c* (Lozano-Urena et al., 2023) which is augmented in *Inpp5e* mutants but decreased in E11.5 *Gli3* conditional mutants (Supp. Table 1). Thus, the effects of Igf signalling on neural progenitor behaviour appear developmentally regulated and require further investigations.

In summary, creating a fully functional cerebral cortex heavily relies on precise cell-cell communication and thus primary cilia. Most notably, these tiny organelles are essential for producing Gli3R which is critical not only for suppressing Sonic hedgehog signalling to prevent a ventralisation of the developing cortex (Kuschel et al., 2003; Tole et al., 2000) but also for controlling the timing of neuronal differentiation in a Shh independent manner (Hasenpusch-Theil et al., 2018). Rescue experiments involving the reintroduction of Gli3R have underscored this important function and achieved remarkable recoveries in restoring cortical neurogenesis (Hasenpusch-Theil et al., 2020), olfactory bulb formation (Besse et al., 2011) and corpus callosum development (Laclef et al., 2015; Putoux et al., 2019). The genes, however, that act downstream of cilia and are directly regulated by *Gli3* remained largely elusive. Our findings address this knowledge gap and provide a detailed list of candidate target genes highlighting two novel potential pathways for further exploration. Thereby, we shed light on the mechanisms by which cilia orchestrate aspects of cortical development and contribute to a more comprehensive apprehension of ciliary functions.

## MATERIAL AND METHODS

### Mice

All experimental work was carried out in accordance with the UK Animals (Scientific Procedures) Act 1986 and UK Home Office guidelines. All protocols were reviewed and approved by the named veterinary surgeons of the College of Medicine and Veterinary Medicine, the University of Edinburgh, prior to the commencement of experimental work. *Inpp5e*^Δ^, *Gli3* conditional (*Gli3*^fl^) mouse mutants and the *Emx1Cre* driver line have been described previously (Blaess et al., 2008; Gorski et al., 2002; Jacoby et al., 2009). *Inpp5e*^Δ/+^ mice were interbred to generate *Inpp5e*^Δ/Δ^ embryos; exencephalic *Inpp5e*^Δ/Δ^ embryos which made up ca. 25% of homozygous mutant embryos were excluded from the analyses. Wild-type and *Inpp5e*^Δ/+^ litter mate embryos served as controls. To generate Gli3 conditional mutants, *Emx1Cre*;*Gli3*^fl/+^ mice were interbred with *Gli3*^fl/+^ animals; *Gli3*^*flox/flox*^, *Gli3*^*flox/+*^,*Emx1Cre* and *Gli3*^*flox/+*^ embryos served as controls. Embryonic (E) day 0.5 was assumed to start at midday of the day of vaginal plug discovery. Transgenic animals and embryos from both sexes were genotyped as described (Hasenpusch-Theil et al., 2012; Jacoby et al., 2009). For each marker and each stage, 3-8 embryos were analysed.

### In situ hybridisation, immunohistochemistry and X-Gal staining on sectioned embryonic brains

In situ hybridisation on 12 μm coronal paraffin sections of E12.5 mouse brains were performed as described previously (Theil, 2005). Digoxigenin-labelled antisense probes were generated from the following cDNA clones: *Pappa* (Genepaint riboprobe T37932), *Sall3* (Genepaint riboprobe T38908). For the reporter gene analysis of in utero electroporated embryos, brains were dissected in PBS and fixed for 3 hours in 4% PFA. After embedding in OCT/sucrose, 14 μm coronal cryosections were analysed by immunofluorescence using an antibody against GFP (1:1000; Abcam), followed by a nuclear counterstain with TO-PRO-1 (1:2000, Invitrogen) as described previously (Hasenpusch-Theil et al., 2017). Adjacent sections were stained between 3 and 24 hours with X-Gal at 37ºC and counterstained with Fast RED (Hasenpusch-Theil et al., 2017).

### Plasmid construction and mutagenesis

All genomic DNA fragments were generated via PCR using wild-type genomic DNA (for oligonucleotides see Table S2). Enhancer sequences were subcloned using a TOPO TA cloning kit (Invitrogen) and verified by sequencing. Putative Gli3 binding sites were mutated using the QuickChange Site-Directed Mutagenesis Kit (Stratagene) (for oligonucleotides used in mutagenesis see Table S3). All mutations were confirmed by sequencing. To test for enhancer activity, wild-type and mutant regulatory elements were subcloned into the *lacZ* reporter gene vector pGZ40 upstream of a human *β-globin* minimal promoter (Yee and Rigby, 1993).

### Electrophoretic mobility shift assay

Electrophoretic mobility assays were performed with biotin labelled oligonucleotides using purified GST and GST-Gli3 fusion protein as described previously (Hasenpusch-Theil et al., 2017). The binding reactions were separated on native 5% acrylamide gels and transferred onto positively charged nylon membranes (Roche) with a Perfect Blue Semi-dry electro blotter (60 minutes at 120volts, 5mA). After UV crosslinking, biotin labelled probes were detected using a Chemiluminenscent Nucleic Acid Detection Module (Thermo Scientific #89880) according to manufacturer’s instructions and imaged using a Kodak BioMaxXAR film.

For oligonucleotide sequences covering the wild-type or mutated Gli3 binding sites see Table S4. The exchanged nucleotides in the mutated forms are underlined. Wild-type, and Gli3 binding site mutant oligonucleotides were used as specific and unspecific competitors, respectively, in a 10- to 100-fold molar excess.

### In utero electroporation

E13.5 pregnant mice were anesthetized with isoflurane and the uterine horns were exposed. *LacZ* reporter gene plasmids and a *GFP* expression plasmid were co-injected into the lateral ventricle at 1mg/ml each with a glass micropipette. The embryo in the uterus was placed between CUY650 tweezer-type electrodes (Nepagene). A CUY21E electroporator (Nepagene) was used to deliver six pulses (30 V, 50 ms each) at intervals of 950 ms. The uterine horns were placed back into the abdominal cavity and embryos were allowed to develop for 24 hours before further processing for immunofluorescence. For each construct, at least 3 different embryos were analysed.

### Bulk mRNA-seq and Bioinformatic Analyses

For bulk mRNA-seq experiments, dorsal telencephalic tissue was dissected from E12.5 *Inpp5e* mutant embryos to generate four different replicates per genotype (control: *Inpp5e*^+/+^ and *Inpp5e*^Δ/+^; mutant: *Inpp5e*^Δ/Δ^). Total RNA was extracted using RNeasy Plus Mini Kit (Qiagen). After assessing the integrity of the RNA samples with an Agilent 2100 Bioanalyzer, (RIN > 8), all RNAs were further processed for RNA library preparation and sequenced (paired-end, 50bp reads) on an Illumina NovaSeq platform at Edinburgh Genomics (University of Edinburgh). Sequencing quality was checked using FastQC software and reads were aligned to the *Mus musculus* reference genome (genome assembly mm10) and analysed using STAR alignment software. The featureCounts tool (Liao et al., 2014) was used to quantify gene expression. Count normalization and differential gene expression analyses were conducted in RStudio using the DESeq2 package (Love et al., 2014). Principal component analyses and hierarchical clustering were applied to normalized count data. Genes were annotated the biomaRt software package (Durinck et al., 2009). Differentially expressed genes were selected based on an adjusted p-value <0.05 and are summarized in Table S1. Gene ontology analyses was performed using Clusterprofiler Software (Wu et al., 2021) in the annotation category BP. Strongly enriched terms had a score of <0.05 after Benjamini-Hochberg multiple test correction.

## ACKNOWLEDGEMENTS

We are grateful to Drs Thomas Becker and John Mason for critical comments on the manuscript, and Stéphane Schurmans for the *Inpp5e*^Δ/+^ mouse line.

## COMPETING INTERESTS

The authors declare no competing or financial interests

## FUNDING

This work was supported by grants from the Biotechnology and Biological Sciences Research Council (BB/P00122X/1) and from the Simons Initiative for the Developing Brain (SFARI -529085) to TT.

## DATA AVAILABILITY

Raw data from gene expression profiling were submitted to the European Nucleotide Archive (ENA) under accession numbers E-MTAB-14015.

## DIVERSITY AND INCLUSION STATEMENT

One or more of the authors of this paper self-identifies as a member of the LGBTQ+ community.

## Notes

### Competing Interest Statement

The authors have declared no competing interest.

