## Supplementary Figures for "Identification of *Pappa* and *Sall3* as Gli3 direct target genes acting downstream of cilia signalling in corticogenesis"

**Supplementary Figure 1**

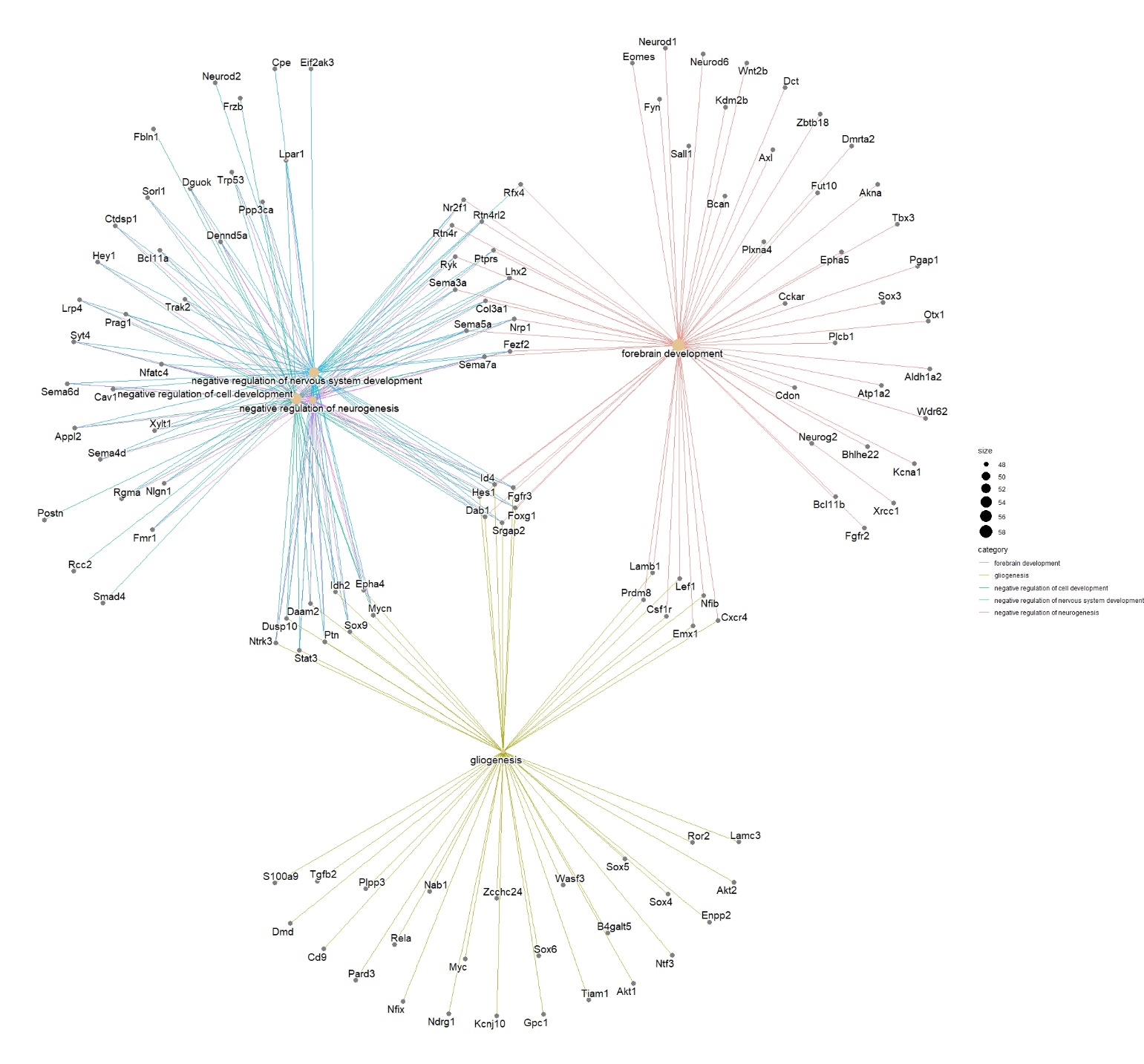

**Supplementary Figure 1: Network plot of down-regulated genes**.

**Supplementary Figure 2**

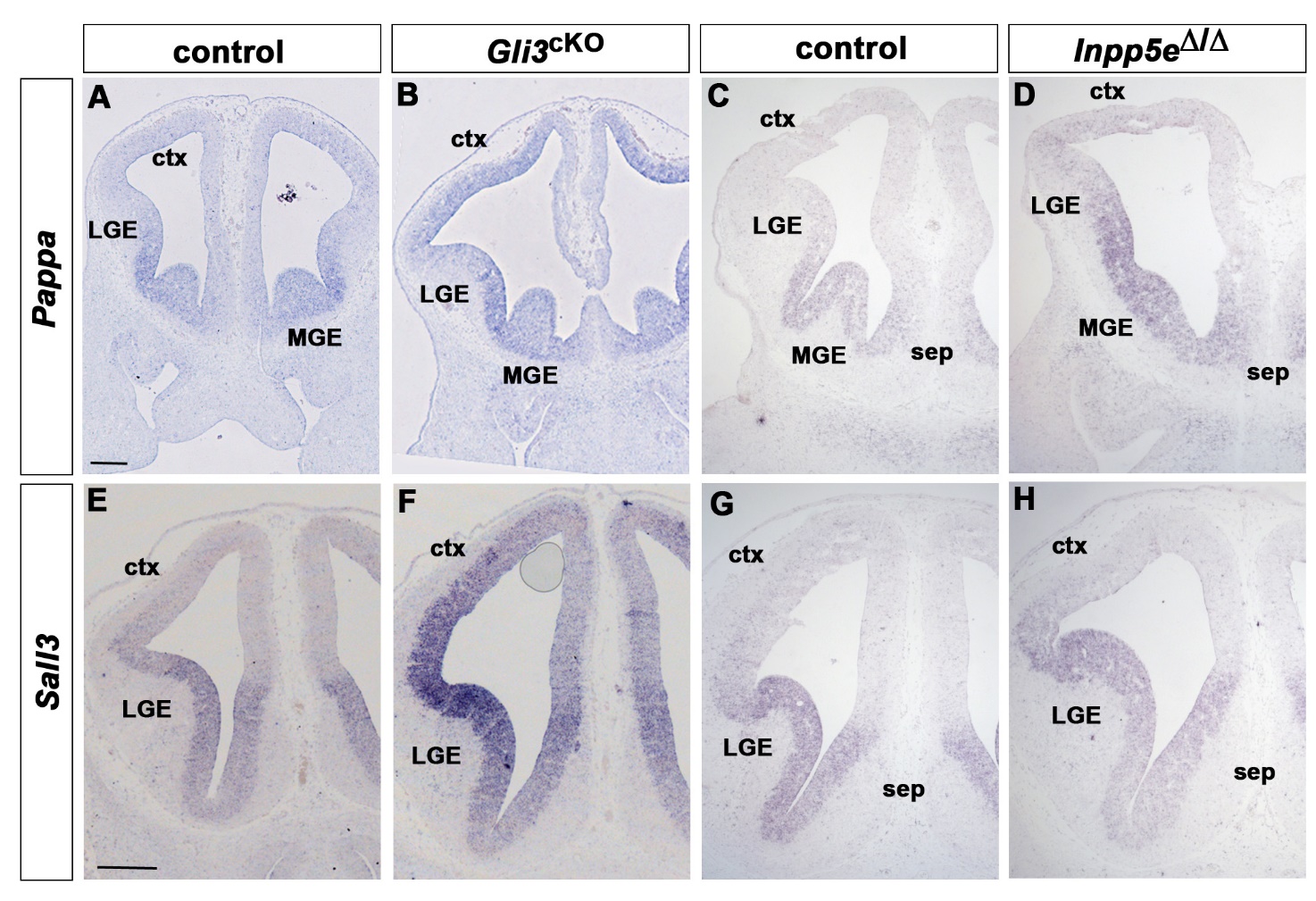

**Supplementary Figure 2: *Pappa* and *Sall3* expression in the *Gli3*^cKO^ and *Inpp5e*^Δ/Δ^ mutant telencephalon**. Coronal sections of the E12.5 forebrain of the indicated genotypes were in situ hybridised with the indicated probes. (A, C) *Pappa* expression is confined to the septum (sep), medial ganglionic eminence (MGE) and the ventral most lateral ganglionic eminence (LGE) of control embryos. (B, D) Ectopic *Pappa* expression throughout the cortex of *Gli3*^cKO^ embryos and in the dorsolateral telencephalon of *Inpp5e* mutant embryos. (E, G) *Sall3* expression is confined to the ventral telencephalon in control embryos. (F, H) *Gli3*^cKO^ embryos display ectopic *Sall3* transcripts in the developing cortex while *Inpp5e* mutants present with a more restricted up-regulation in the dorsolateral telencephalon. Scale bars: 200 μm.

**Supplementary Figure 3**

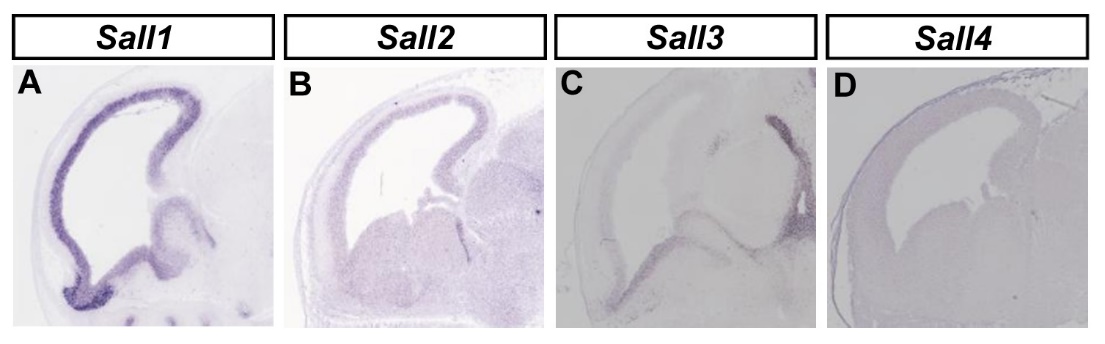

**Supplementary Figure 3: *Sall* gene expression in the E14.5 mouse cortex**. Images were taken from Genepaint.

**Supplementary Figure 4**

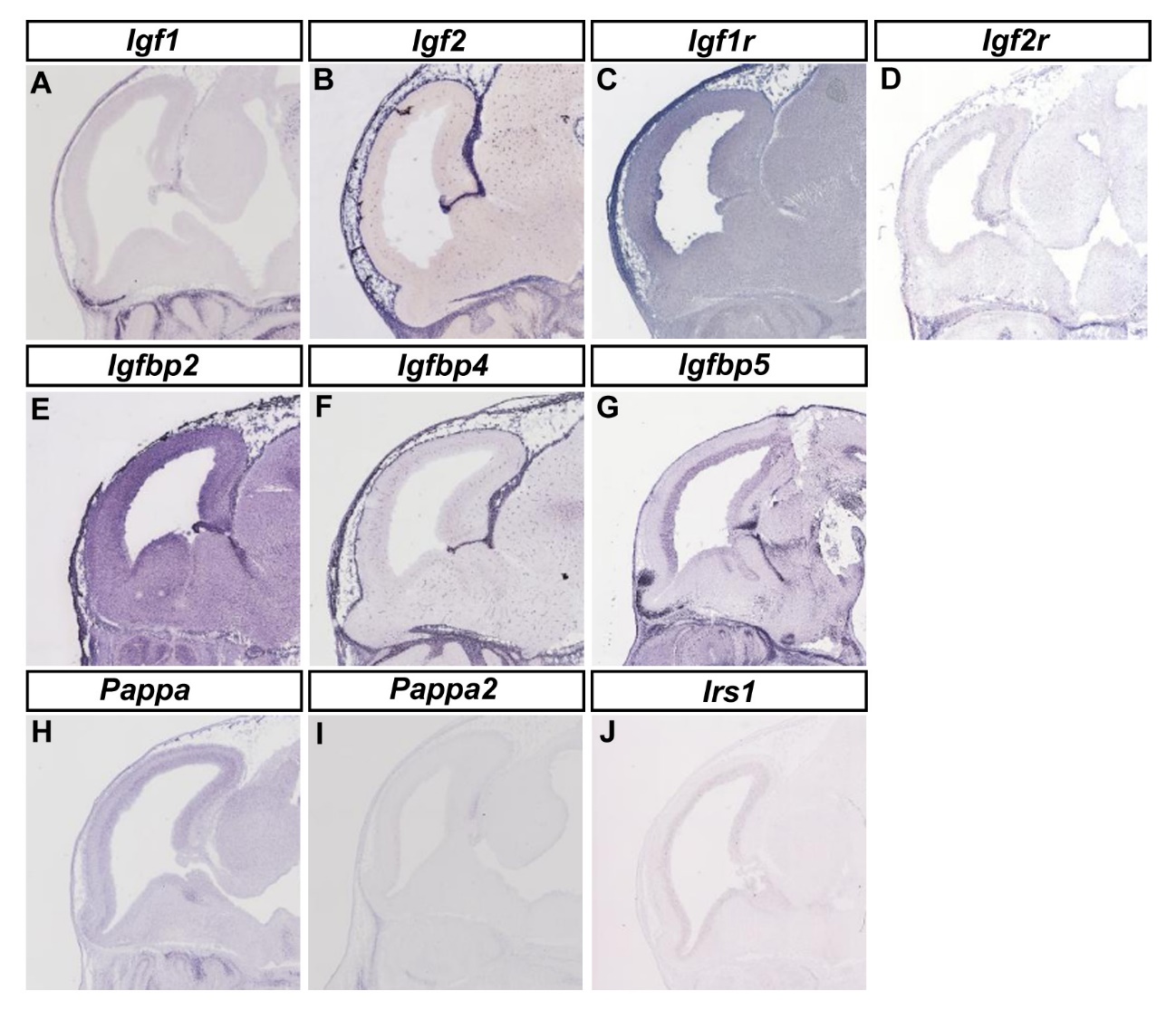

**Supplementary Figure 4: Expression of genes encoding Igf signalling components in the E14.5 mouse cortex**. Images were taken from Genepaint.
