## Supplementary Tables 2-4 for "Identification of *Pappa* and *Sall3* as Gli3 direct target genes acting downstream of cilia signalling in corticogenesis"

**Supplementary Table 2**

**Cloning *Pappa* enhancer**

Forward primer	AAAGGTACCTCTCTGCTACCATCCGTCCT

Reverse primer	AAAGTCGACAGGCCTGCAGAAGATAAGGG

**Cloning *Sall3* enhancer**

Forward primer	AAAAAGCTTGCATCTCAAGTCGGACGAGG

Reverse primer	AAACTCGAGATAGCGACCGTTCTGCACTT

**Supplementary Table 3**

**Oligonucleotides used for mutagenesis**

*Pappa* binding site 1

Forward primer	GCCATCTCGGGGGT**AT**CCCTCCGCTCTTTC

Reverse primer	GAAAGAGCGGAGGG**AT**ACCCCCGAGATGGC

*Pappa* binding site 2

Forward primer	GAGCCATAGATGAGGGT**AT**CCATTTGGGGGAGG

Reverse primer	CCTCCCCCAAATGG**AT**ACCCTCATCTATGGCTC

*Sall3* binding site 1

Forward primer	GTTCTGGGGCGC**AT**ACCCAGGTCGGGAAGC

Reverse primer	GCTTCCCGACCTGGGT**AT**GCGCCCCAGAAC

*Sall3* binding site 2

Forward primer	GGGGACCTCAGGGT**AT**CCCAGAGTTCTGGG

Reverse primer	CCCAGAACTCTGGG**AT**ACCCTGAGGTCCCC

**Supplementary Table 4**

**Oligonucleotides used for EMSA**

*Pappa* Binding site 1

Biotinylated forward primer	Btn-CATCTCGGGGGTGGCCCTC

Forward primer	CATCTCGGGGGTGGCCCTC

Reverse primer	GAGGGCCACCCCCGAGATG

Forward mutated primer	CATCTCGGGGGT**AT**CCCTC

Reverse mutated primer	GAGGG**AT**ACCCCCGAGATG

*Pappa* Binding site 2

Biotinylated forward primer	Btn-CATAGATGAGGGTGGCCATTTG

Forward primer	CATAGATGAGGGTGGCCATTTG

Reverse primer	CAAATGGCCACCCTCATCTATG

Forward mutated primer	CATAGATGAGGGT**AT**CCATTTG

Reverse mutated primer	CAAATGG**AT**ACCCTCATCTATG

*Sall3* Binding site 1

Biotinylated reverse primer	CGACCTGGGTGGGCGCC-Btn

Forward primer	GGCGCCCACCCAGGTCG

Reverse primer	CGACCTGGGTGGGCGCC

Forward mutated primer	GGCGC**TA**ACCCAGGTCG

Reverse mutated primer	CGACCTGGGT**TA**GCGCC

*Sall3* Binding site 2

Biotinylated reverse primer	CTCTGGGCCACCCTGAGGTC-Btn

Forward primer	GACCTCAGGGTGGCCCAGAG

Reverse primer	CTCTGGGCCACCCTGAGGTC

Forward mutated primer	GACCTCAGGGT**AT**CCCAGAG

Reverse mutated primer	CTCTGGG**AT**ACCCTGAGGTC
